# Ramping Dynamics in the Frontal Cortex Unfold Over Multiple Timescales During Motor Planning

**DOI:** 10.1101/2024.02.05.578819

**Authors:** R.O. Affan, I.M. Bright, L.N. Pemberton, N.A. Cruzado, B.B. Scott, M.W. Howard

**Affiliations:** Graduate Program in Neuroscience, Boston University, Boston, MA; Department of Psychological and Brain Sciences, Boston University, Boston, MA 02215

## Abstract

Plans are formulated and refined over the period leading to their execution, ensuring that the appropriate behavior is enacted at just the right time. While existing evidence suggests that memory circuits convey the passage of time through diverse neuronal responses, it remains unclear whether the neural circuits involved in planning behavior exhibit analogous temporal dynamics. Using publicly available data, we analyzed how activity in the frontal motor cortex evolves during motor planning. Individual neurons exhibited diverse ramping activity throughout a delay interval that preceded a planned movement. The collective activity of these neurons was useful for making temporal predictions that became increasingly precise as the movement time approached. This temporal diversity gave rise to a spectrum of encoding patterns, ranging from stable to dynamic representations of the upcoming movement. Our results indicate that neural activity unfolds over multiple timescales during motor planning, suggesting a shared mechanism in the brain for processing temporal information related to both past memories and future plans.

## Introduction

Adaptive behavior necessitates executing appropriate actions precisely when needed. Humans and other animals are capable of planning in the present for future actions. Effective motor planning demands ongoing adjustments in response to evolving situations and a steadily increasing commitment as the moment for action execution draws near. Such computational considerations hint at an underlying neural system with dynamics inherently capturing the ongoing flow of time.

The task of representing and updating the timing for future planned events parallels the challenge of estimating the timing of past occurrences. Hippocampal ’time cells’ activate in sequences following task-related events, forming distinct patterns for various events ^1,2,3^. Conversely, ’temporal context cells’ in the entorhinal cortex exhibit rapid increases or decreases in activity upon encountering an event, subsequently returning exponentially to baseline ^4,5^, with each neuron displaying a unique relaxation time constant. Whereas time cells create continuous timelines by sequencing activation through intervals, temporal context cells achieve this by spanning a range of time constants. It remains uncertain whether these temporal dynamics are exclusive to brain circuits associated with memory or if they extend more broadly across the cortex for future action planning.

The anterior lateral motor (ALM) cortex plays a pivotal role in motor planning for mice. Within the ALM, certain neurons demonstrate a gradual ramping up or down in firing rates during a delay interval that bridges sensory cues and the ensuing motor response. Prior research has typically viewed ramping as a population-level characteristic of ALM activity, distinct from how the animal’s impending movement direction is encoded. In our study, we delve into the variability in ramping dynamics and direction selectivity among ALM neurons. We inquire if ALM cortex ramping activity exhibits a uniform time constant or a spectrum of time constants that span the delay interval leading to action initiation. Utilizing established analytical methods ^5,6^, we determined the ramping time constants of individual ALM neurons from a publicly accessible dataset ^7^. The observed wide array of time constants in neuron ramping activity suggests a universal computational strategy in the brain for integrating past experiences and forthcoming actions within a continuous temporal framework.

## Results

We used a model-based approach to evaluate the temporal modulation of activity in the mouse ALM cortex that was retrieved from an open-source dataset (https://crcns.org/data-sets/motor-cortex/alm-5/about-alm-5). The dataset was originally collected and reported by Inagaki et al. ^7^ and included preprocessed, extracellular recordings from the ALM cortex of mice during an auditory, fixed-delayed response task. In this task, a trial begins with a 1.15 s interval in which mice are presented with a sequence of five 150 ms long 3 kHz or 12 kHz tones (Figure 1a). The final tone is followed by a 2 s long delay period. At the end of this delay period, a 100 ms long 6 kHz tone is presented as a ”Go” cue to instruct mice to lick rightward for the 3 kHz tones or leftward for the 12 kHz tones. Details of the task are described in the original report of the dataset ^7^.

**Figure 1:**
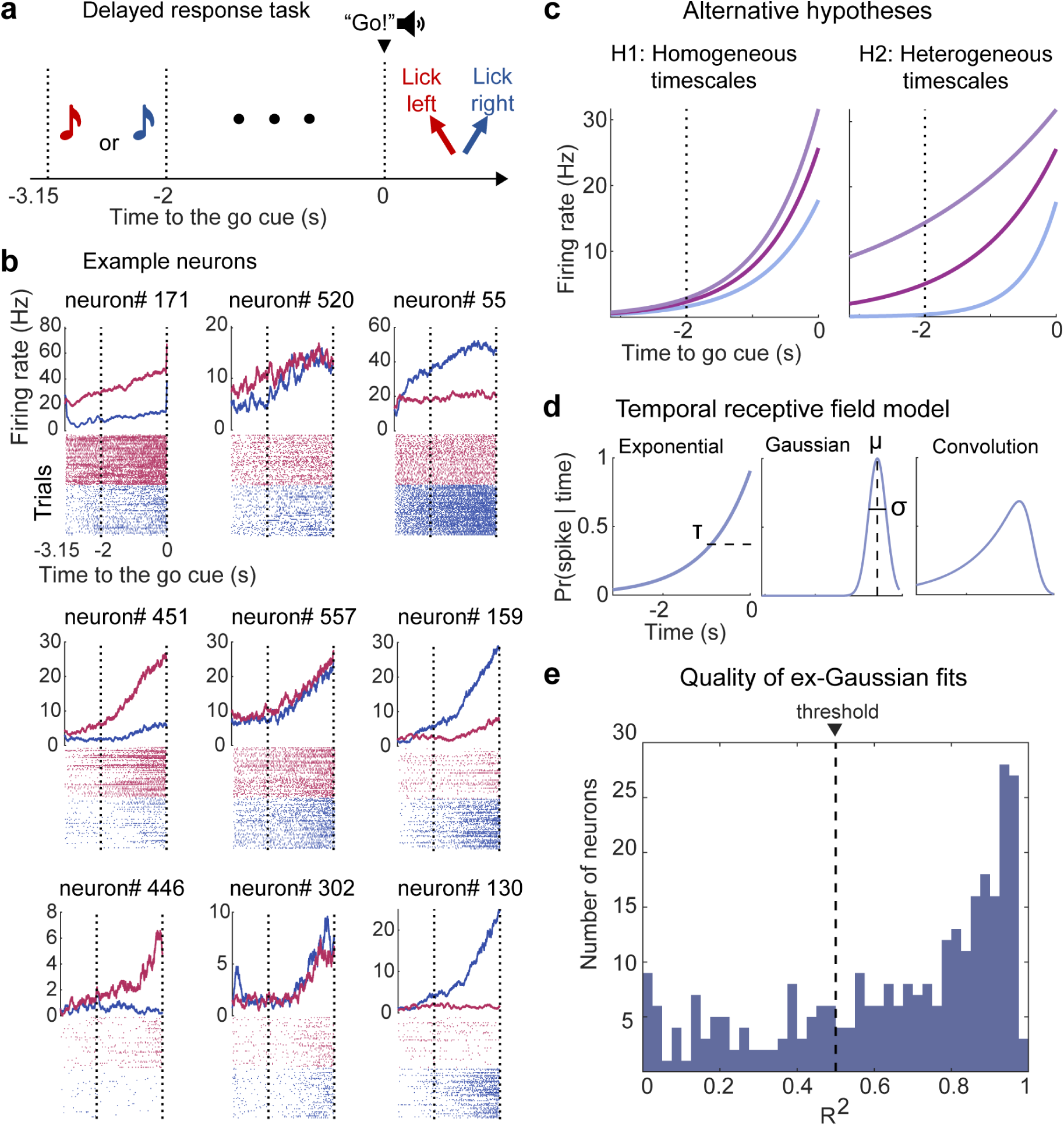
Schematic of the delayed response task and hypotheses for neural activity. **a.** Schematic of the auditory delayed response task from Inagaki et al. ^7^ . Red represents trials where mice are presented with high-pitch tones, instructing them to lick leftward following a Go cue. Blue corresponds to trials with low-pitch tones, instructing them to lick rightward. **b.** Example neurons in the ALM cortex. Top, cross-trial average PSTHs. Bottom, spike rasters from 50 randomly sampled trials for each condition. The colors correspond to the different trial conditions in a. Error trials are excluded from the analyses. Vertical dotted lines mark the start and end of the delay period. Some neurons selectively fire in one type of trial over the other (left column and right column), discriminating the direction of the future lick response. Some neurons gradually ramp up during the delay period (top row), while others ramp up more abruptly at the end of the delay period (bottom row). **c.** Alternative hypotheses for the temporal properties of neural activity during motor planning. H1: The firing rate of individual neurons ramps over the delay period with similar time constants. H2: The firing rate of individual neurons ramps over the period with different time constants. **d.** An ex-Gaussian receptive field model of neuron firing activity. The ex-Gaussian receptive field model (right) is generated from the convolution of an exponential function (right) and a Gaussian function (middle). This model describes the response as a function of time with the time constant parameter *τ* , the peak latency parameter *µ*, and the parameter *σ*. **e.** Quality of the model fit across ramping neurons. Most upward ramping neurons (216 of 302 neurons) were fit by the ex-Gaussian receptive field model with *R*^2^ *>* 0.5 (black dashed line).

As previously reported by Inagaki et al. ^7^, many neurons in the ALM cortex showed firing rates that fluctuate throughout the 2 s long delay period (n = 516 of 667 putative pyramidal units). Some neurons showed an increase in firing rate as time approaches the end of the delay period (n = 302 of 516 neurons), while others showed a decrease from baseline firing (n = 214 of 516 neurons). Most neurons exhibited some level of selectivity in their firing for right- vs left-lick trials (n = 344 of 516 neurons) while others show no preference for one trial type over the other (n = 172 of 516 neurons). The activity of a few example neurons are shown in Figure 1b. The diversity of selectivity and ramping direction in this dataset has been characterized in the original report ^7^. The primary focus of this paper is to evaluate whether motor preparatory ramps, as identified by Inagaki et al. ^7^, have a single homogeneous timescale or if the ramps show heterogeneous timescales across neurons during the delay period (Figure 1c). Neurons that ramp downward are of secondary interest in this paper and are briefly reported in the end of the *Results* section and in the supplement.

### Receptive field model estimates the temporal properties of single-neuron firing

To quantify the temporal characteristics of neural activity in the ALM cortex, we estimated their temporal receptive fields as a convolution of a Gaussian (latency of the activity) and an exponential ramp (Figure 1d). Under this model, the latency of each neuron’s activity is described by the mean of the Gaussian function *µ*. The rate at which each neuron’s activity ramps up or down is described by the time constant of the exponential term *τ* . An additional parameter *σ* determines the standard deviation of the Gaussian function. We classified neurons as either upward or downward ramping, and as selective or non-selective, in accordance with the definitions provided in the original report of this data by Inagaki et al. ^7^. Neurons that show trial-type selectivity were fit with two amplitude terms corresponding to right- and left-lick trials. Neurons that show downward ramping of activity were fit with a negative amplitude term. Details of the parameter estimation procedure are described in *Methods*.

Using this approach, we identified 216 upward ramping neurons in the ALM cortex whose firing activity was fit by the ex-Gaussian receptive field model with an *R*^2^ value greater than 0.5 (Figure 1e). These neurons make up 64.07% of the total 334 putative pyramidal neurons that were fit with *R*^2^ *>* 0.5. Neurons with *R*^2^ values less than 0.5 were excluded from further analysis.

### Neuronal firing in the ALM cortex ramps with heterogeneous time constants

Different ALM neurons displayed firing rates that ramp up at varying speeds over the course of the 2 s delay interval. Slower ramping neurons, such as the three shown at the top row of Figure 2a, began to increase their firing rate as early as the stimulus period (between -3.15 s and -2 s to the Go cue) and gradually reached maximum or minimum firing rate around the Go cue. In contrast, faster ramping neurons, like those shown in the bottom row of Figure 2a, displayed a more abrupt change in firing rate later in the delay period and reached maximum or minimum activity at the time of the Go cue. The variability in ramping dynamics across all 216 ALM neurons is displayed in Figure 2b.

**Figure 2:**
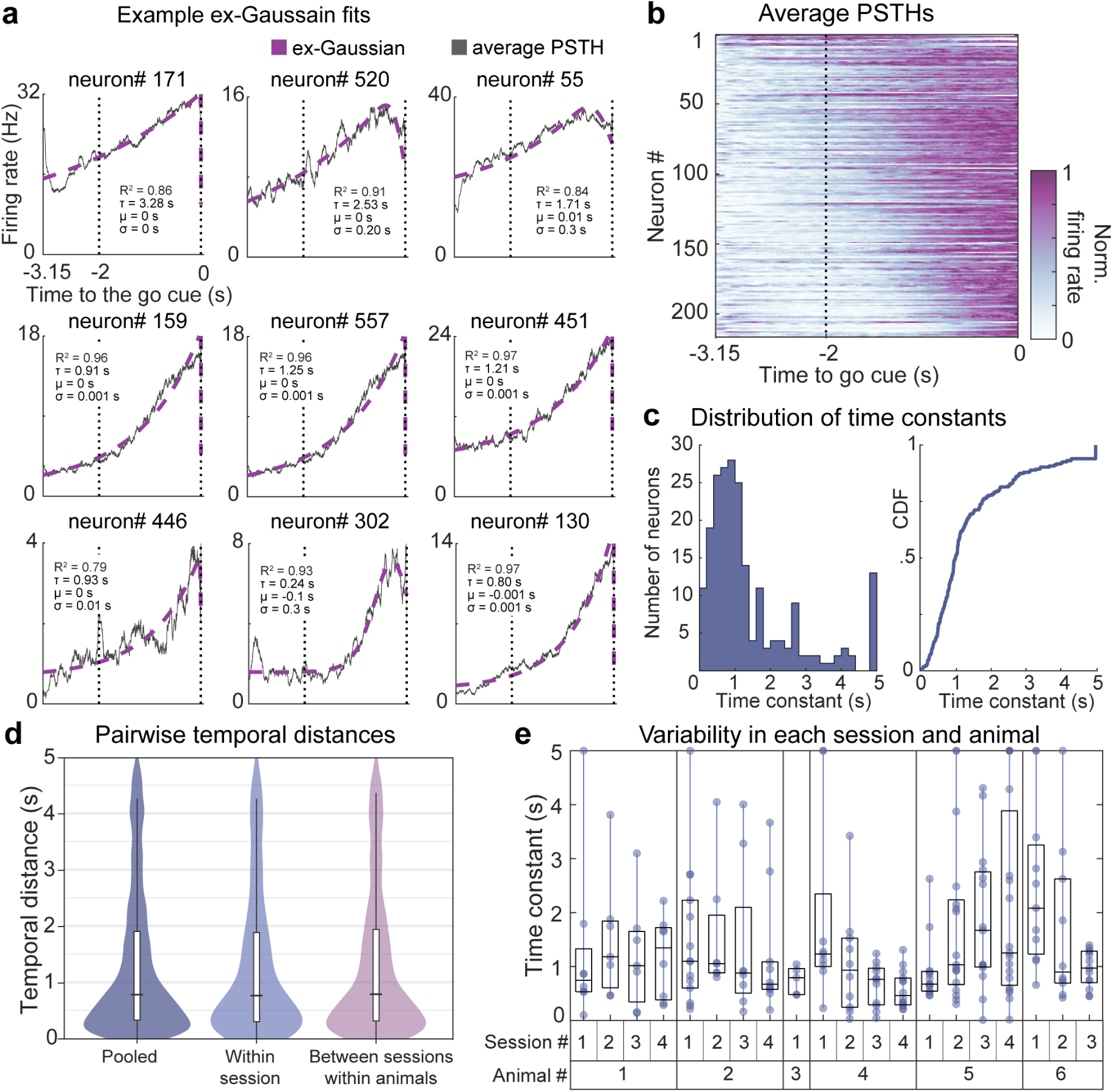
Neuronal firing rates ramp up at various timescales in the ALM cortex. **a.** Ex-Gaussian receptive field model fits for nine example ALM neurons. The ex-Gaussian receptive field model is compared with the averaged firing rate of each neuron 3.15 s preceding the Go cue. Firing rates are depicted as thin black lines. Ex-Gaussian models are depicted as dashed purple lines. These neurons differ in their time constant, estimated as the *τ* of the receptive field models. **b.** Heatmap of normalized firing rates across 216 ALM neurons. Each row represents the averaged normalized firing rate for a single neuron. Neurons were sorted by their estimated value of *τ* , from larger *τ* or longer time constant to smaller *τ* or shorter time constant at the bottom. **c.** Histogram and cumulative distribution of estimated time constants. The histogram on the left shows the marginal distribution of the *τ* values, which describes the time constant of ramping activity for each neuron. The cumulative distribution function (CDF) of *τ* is depicted in the purple solid line on the right side. A few neurons exhibit values of *τ* that hit the upper bound 5 s. **d.** Violin plots show pairwise temporal distances calculated between neurons from all recording sessions and animals (left), the same session (middle), and different sessions of the same animals (right). Temporal distance was measured as the absolute difference between time constant (*τ* ) values. **e.** Box plots representing the variability of time constants (*τ* ) for ALM neurons within individual sessions and across different animals. Estimated *τ* values varied among neurons in each recording session, showing some variability across sessions and minimal variability between animals.

There was a wide range of time constants across neurons, as indicated by a median *τ* of 0.97 s and an interquartile range from 0.57 s to 1.74 s. Notably, 90% of the time constants were less than or equal to 3.42 s. Figure 2c shows the histogram and cumulative distribution of time constant values, revealing a skewed distribution (*skewness* = 1.56) with a heavy-tail (*kurtosis* = 4.64). Indeed, the distribution of time constants closely fits a log-normal (location = *−*0.05 *±* 0.07 (SE) and scale = 0.99 *±* 0.05, LL = -294.47, AIC = 592.94, BIC = 599.69), or gamma (shape = 1.39 *±* 0.12 (SE) and rate = 0.97 *±* 0.10, LL = -285.78, AIC = 575.55, BIC = 582.30) distribution. These patterns indicate that while most neurons exhibit shorter time constants, a greater-than-expected number (under a normal distribution assumption) showed longer time constants. In contrast to the wide range of ramping time constants, there was relatively little variability in peak response times (*µ* = 0 s in 90% of the 216 neurons). Alternative estimates of time constants, inferred from autocorrelation functions ^6^, similarly yielded a wide range of values across these neurons (Figure S1). This pattern of results closely resemble the temporally diverse responses of neurons in the monkey ^5^ and rodent ^4^ entorhinal cortex.

To determine whether the observed variability in time constants reflects heterogeneity among simultaneously recorded neurons, we measured the temporal distances between pairs of neurons in three distinct conditions. In the first condition, pairwise temporal distances were measured between neurons from any session and animal (median = 0.78 s, interquartile range = 0.33 s to 1.90 s). In the second condition, pairwise distances were calculated between neurons in the same session (median = 0.76 s, interquartile range = 0.30 s to 1.88 s). Lastly, pairwise distances were estimated between neurons from different sessions of the same animal in the last condition (median = 0.79 s, interquartile range = 0.31 s to 1.94 s). As shown in Figure 2d, pairwise distances were similar across the pooled, intra-session, and inter-session conditions. This consistency in pairwise distances indicates that the variability of neuronal time constants within a single recording session is as pronounced as the variability across different sessions.

To further dissect the contributions of session-to-session variability and between-animal differences on the heterogeneity of time constants, we employed a generalized linear mixed-effects model with the random effects of sessions nested within animals. Variance component analysis revealed that variability across sessions accounted for approximately 9.39% of the variance in neuronal time constants (95% CI = [4.09%, 16.01%], *p < .*001 nonparametric bootstrap), while variability across animals captured around 0.84% of the variance (95% CI = [0%, 5.04%], (*p* = 0.302). Figure 2d shows the distribution of time constants across individual sessions and animals. Notably, a considerable proportion of the variance remained after accounting for inter-session and inter-animal variability (89.77%, 95% CI = [83.45%, 94.68%], (*p < .*001). Coupled with the pairwise distances observed within the same sessions, this result suggests that the heterogeneity in time constants may reflect temporal diversity within a population of simultaneously recorded neurons in the ALM cortex.

### Heterogeneous ramping dynamics allows time to be encoded at different scales

Our analyses reveal a broad range of time constants among individual ALM neurons, therefore we asked whether this heterogeneity may have implications on dynamics at the population level. Given the proposal that ramping activity in the ALM may reflect time estimation related to urgency signals ^8^, we first evaluated whether the joint activity of the 216 neurons carries latent information about time. We trained a decoder using linear discriminant analysis (LDA) to estimate time throughout the 2 s delay period using the population activity of the 216 neurons. We measured the confidence of the decoder at each 100 ms time bins during the delay interval as the log posterior across the range of possible time estimates. The decoder made predictions that were proximal to the actual time bins with an average error of 0.19 s, resulting in a diagonal band across the decoding matrix in Figure 3a. A *χ*^2^ goodness of fit test indicated that this pattern was different from a uniform distribution that would result from decoding at chance level, *χ*^2^(992) = 2.142 and *p < .*001. A linear regression of the average decoding error at each time bin yielded a negative slope of 0.057 *±* 0.008 (Figure 3b). These results indicate that the population activity of these ALM neurons conveys information that can be used to decode time with increasing precision as time approaches the Go cue.

**Figure 3:**
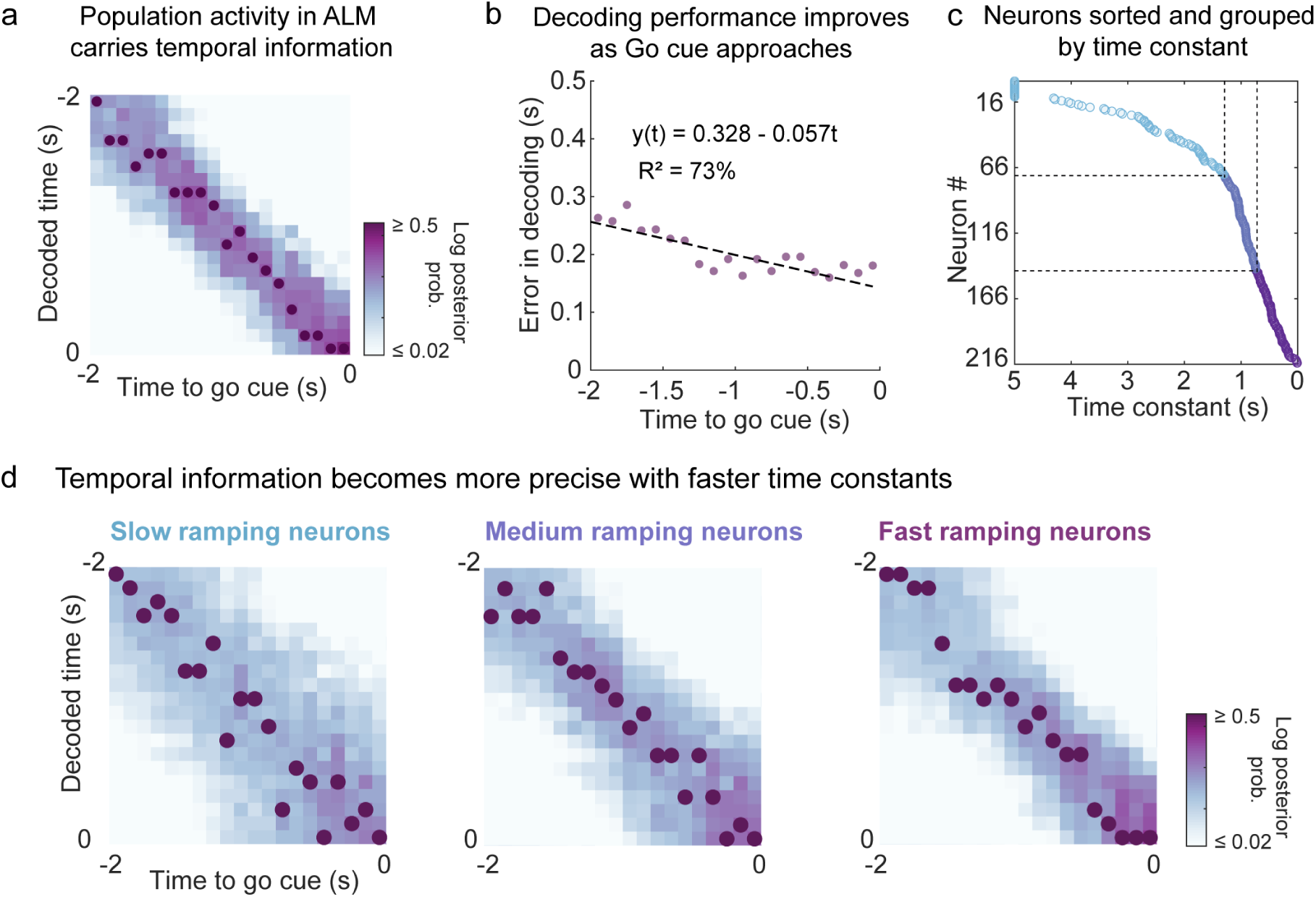
Population activity in the ALM cortex conveys information about time relative to the Go cue. **a.** A linear discrimination analysis (LDA) decoder was trained to classify 100 ms time bins along the 2 s delay interval on the y-axis from the population activity of ramping neurons at different 100 ms time bins along the 2 s interval on the x-axis. The color gradient represents the log posterior probability (confidence) of the predictions. Dark purple dots indicate the positions of maximum log posterior probability for along the y-axis for each time bin on the x-axis. **b.** Decoding error as a function of time within the delay intervals. The x-axis shows the 100 ms time bins that tile the 2 s long intervals preceding the Go cue. The y-axis shows the average error between the decoder’s predictions and the actual time in seconds. A linear fit of the decoding error is plotted as a dashed line, showing a decrease of decoding error as time reaches the end of the delay interval. **c.** ALM neurons sorted and grouped by *τ* . The 216 ramping neurons on the y-axis were sorted by their estimated time constant *τ* on the x-axis. Neurons were categorized into three equal groups (n = 72 each), labeled as fast (dark purple dots), medium (light purple dots), and slow (light blue dots) based on their time constants. The values of *τ* on the boundaries marked by the dashed lines are 0.74 s and 1.29 s **d.** Time decoded from fast, medium, and slow ramping neurons. Three LDA decoders (a) were exclusively trained on the population activity of neurons in the slow, medium, and fast group (c). The slow ramping population yielded a light broad diagonal (right). The fast ramping population yielded a dark narrow diagonal (left).

Next, we investigated whether the temporal information present in the population activity is dependent on the specific range of time constants among its constituent neurons. Neurons were sorted by their estimated *τ* value then categorized them into slow-, medium-, and fast-ramping groups (Figure 3c), each consisting of 72 of the 216 neurons. Neurons in the slow-ramping group have*τ* values greater than 1.29 s, while neurons in the fast-ramping group have *τ* values less than 0.74 s. LDA analysis was performed to decode time using the population activity of each group separately, as shown in Figure 3d. The precision of temporal estimation differed between the three groups during 1 s preceding the go cue (*F* (2, 27) = 9.48, *p < .*001). Not surprisingly, temporal information was less precise among neurons with the slowest time constant (average decoding error = 0.378 s, SD = 0.036 s) than among the medium-ramping (average decoding error = 0.264 s, SD = 0.030 s) and fast-ramping neurons (average decoding error = 0.278 s, SD = 0.101 s), all *p*s *< .*001. The difference in temporal resolutions as a function of ramping time constant is demonstrated in Figure 3d as the diagonal of the LDA matrix becomes more narrow and dark with faster time constants. Even with this difference, temporal information can be successfully decoded well above chance for all three subpopulations (*χ*^2^(992) *>* 63, 137 and all *p*s *< .*001). The results demonstrate that temporal information is present in the population activity of ALM cortex across a wide range of timescales, corresponding to the ramping time constants of its constituent neurons.

### Temporal heterogeneity affects the encoding of future movement direction

The timescale with which neural activity fluctuates can determine how it contributes to different functions. For instance, neurons with longer time constants have previously been shown to be more involved in maintaining items in working memory compared to those with shorter time constants ^9,10^. Thus, we evaluated how the temporal heterogeneity relates to the encoding of upcoming movements in the ALM cortex. We first examined the joint distribution of time constants and trial-type selectivity across individual ALM neurons. Selectivity was measured as the normalized difference between the average firing rates estimated at the delay interval for correct right-lick trials and correct left-lick trials, as previously defined in the original report of this dataset ^7^. Figure 4a shows the marginal distribution of normalized selectivity during the delay interval. As previously reported by Inagaki et al. ^7^, ALM neurons show a continuous distribution of selectivity with a contralateral bias or right-lick preference (median *selectivity* = 0.03, interquartile range = -0.14 to 0.30).

**Figure 4:**
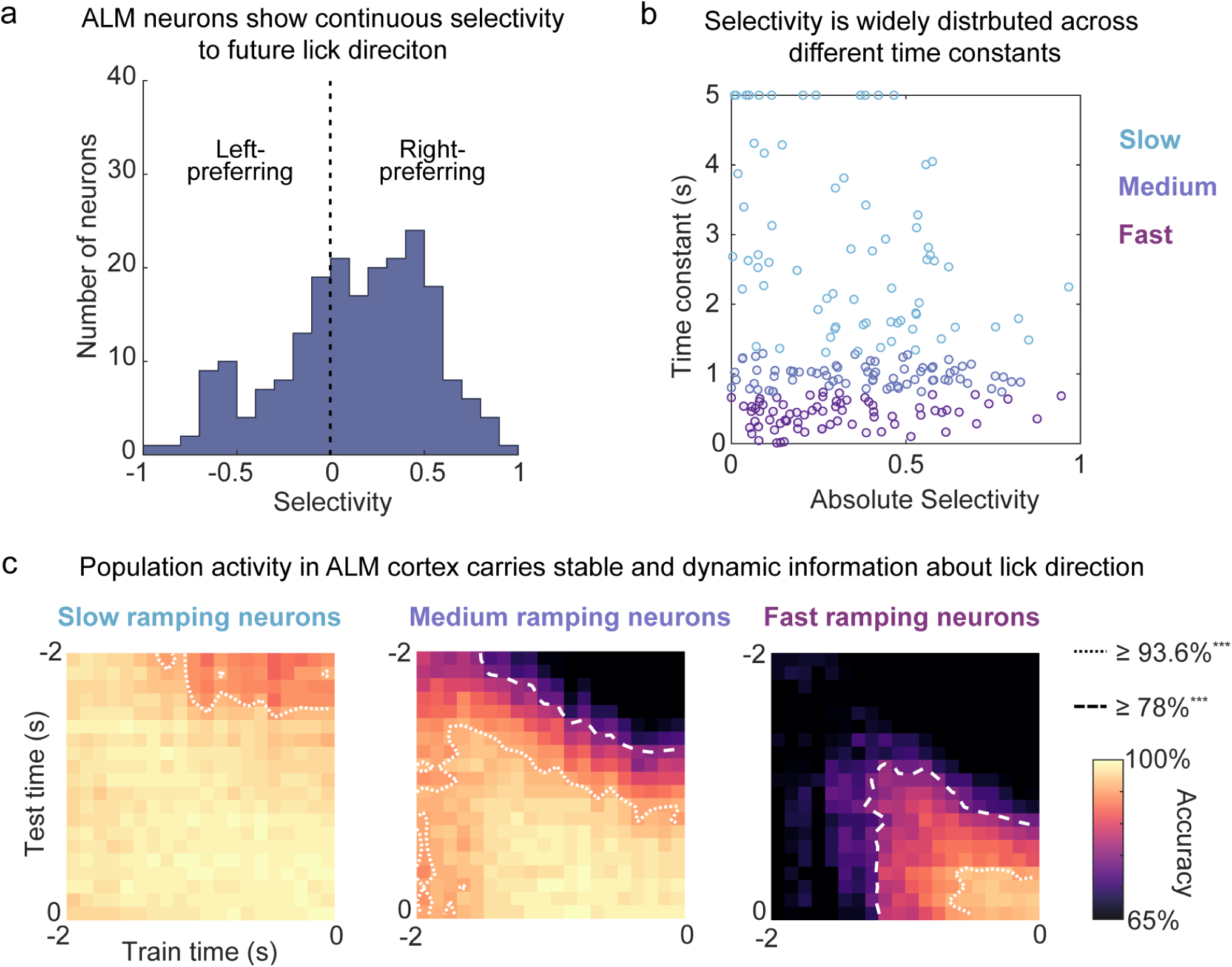
Population activity in the ALM carries stable and dynamic information about trial types. **a.** Trial type selectivity is distributed across ALM neurons. The histogram of normalized selectivity value across the 334 neurons. Positive selectivity values correspond to higher firing rates across the right-lick trials relative to the leftlick trials. Negative selectivity values correspond to more activity in left-lick over right-lick trials. Selectivity was constrained to activity during the delay period only. Only correct trials were used. **b.** Scatterplot of selectivity and time constants. Information about trial type (i.e., low-pitch/lick-right vs high-pitch/lick-left) was distributed broadly across time constants. The y-axis shows the absolute value of the normalized selectivity index form panel a (see text for details). The x-axis shows the estimated (*τ* ) values. Each dot represents an individual neuron colored by the group they belong to (see Figure 3c). Slow-ramping neurons are light blue dots. Fast-ramping neurons are dark purple dots. Medium-ramping neurons are light purple dots. **c.** Accuracy matrices for the cross-temporal decoding (CTD) of trial types using the activity of slow-, medium-, and fast-ramping neurons. LDA classifiers were trained to predict whether animals correctly licked left or right in each trial, at every possible pair of 100 ms time bins along the 2 s delay interval. Each CTC was exclusively trained on the population activity of the slow, medium, or fast ramping neurons. The color gradient corresponds to decoding accuracy (%). The outermost contour line in each matrix indicates where classifiers performed above chance (*p < .*001, one proportion z-test). The interior contour line highlights further significant increases at *p < .*001 using two proportion z-tests. All three groups yielded decoding performance above chance. Decoding performance for the slow-ramping neurons was consistently high throughout the entire span of the delay interval. Decoding performance for the-fast ramping neurons increased smoothly and peaked as time approaches the Go cue.

A scatterplot of *τ* as a function of the absolute value of selectivity is shown in Figure 4b for neurons in the slow-, medium-, and fast-ramping group. There was no significant correlation between *τ* and absolute selectivity value (Kendall’s *τ* = 0.02, *p* = 0.82). To evaluate the strength of this null finding, we calculated a Bayes factor (*BF*_01_ = 1.72) for whether neurons with varying time constants exhibit similar levels of trial-type selectivity (null hypothesis) or differ in their selectivity (alternative hypothesis). This value indicates that the data are 1.72 times more likely under the null hypothesis, which suggests that information about the direction of upcoming movement is conveyed in the ALM across a range of timescales, without a strong dependence on the neurons’ time constants.

Next, we assessed how neurons with different time constants contribute to trial-type encoding at the population level. To this end, we employed cross-temporal decoding (CTD) to predict the direction of future movement from neural population activity when decoders are trained and tested on all possible pairs of time points within the delay period ^11,12,10,13^. The results of CTD are displayed in Figure 4c as 2D accuracy matrices for the slow-, medium-, and fast-ramping groups. Decoding performance for identical training and testing times are depicted in the diagonal of each matrix. These diagonal elements estimate the presence of information in neuronal responses as a function of time. The off-diagonal elements represent accuracy when training and testing was performed in different time points during the delay, thus measuring the similarity in the neural representation of information across different time points.

High accuracy was observed along the diagonal of the accuracy matrix for neurons in the slowramping group. This demonstrates that trial-type information is sustained in the population activity of these neurons throughout the entire delay period. In contrast, decoding accuracy for the fast-ramping group was comparatively lower during the early half of the delay, then increased to match the performance levels observed in the slow-ramping group. These results indicate that neurons with different time constants contribute to trial-type selective signals at the population level that emerge with varying latency over the delay period.

Decoding accuracy was uniformly high across the off-diagonal elements for the slow-ramping group, demonstrating a stable representation ^14^ of trial type that generalizes across different time points in the delay period. Alternatively, off-diagonal accuracy decreased in the the fast-ramping group as decoders were trained and tested in distal time points (contours lines in Figure 4c). This decrease in performance implies a dynamic representation ^15^, whereby activity patterns that distinguish trial conditions gradually evolve at the population level as time passes during the delay. The contrast in decoding patterns between slow- and fast-ramping neurons suggests that temporal heterogeneity might impact how information about the upcoming movement of the animal might be conveyed to downstream neurons.

### Downward ramping neurons

As previously reported by Inagaki et al. ^7^, some neurons showed a downward ramping of activity from baseline firing rate (n = 214 of the 516 delay-active pyramidal neurons). The firing rates of downward ramping neurons were poorly fitted by the ex-Gaussian model (Mean *R*^2^ *±* SD = 0.49 *±* 0.25). Only 55.14% of the downward ramping neurons had an *R*^2^ value greater than 0.5, whereas 71.52% of the upward ramping neurons surpassed this criterion. Furthermore, LDA analyses revealed that the population activity of downward ramping neurons yielded worse performance in decoding time (average decoding error = 0.264 s, SD = 0.030 s) and trial types (Accuracy range = [ 50.81% - 72.58%]) than the population activity of upward ramping ones. The findings pertaining to downward ramping neurons are plotted in Figure S2.

## Discussion

Neural activity in the ALM cortex is heterogeneous during motor planning, but the nature of this heterogeneity has remained largely unexplored. Here, we reveal a diversity of ramping time constants among ALM neurons in a dataset made publicly available by Inagaki et al. ^7^. Neurons ramp at different speeds throughout a 2 s delay before a motor response, reflecting the temporal heterogeneity among ALM neurons (Figure 1 & Figure 2). This variability allows time to be decoded over a wide range of temporal resolutions from population activity (Figure 3). Because some ramping neurons show selective responses to the direction of the animal’s upcoming movement, movement direction can be decoded from the population at different onsets during the delay and with varying degrees of stability over time (Figure 4). These results resemble previous observations in other brain areas involved in encoding or maintaining memory, such as the entorhinal ^4,5^ and prefrontal cortex ^14,16^. Thus, our findings suggest that temporally diverse ramping dynamics reflect a general mechanism in the cortex that can implicitly represent the passage of time with respect to the occurrence of a past event or the anticipation of an upcoming one.

Previously, Finkelstein et al. ^8^ demonstrated that training an artificial neural network to accurately mimic the diverse responses of ALM neurons requires a recurrent structure and a ramping input. Further work is needed to identify the specific pattern of recurrent and input connections that contribute to the emergence of heterogeneity in this network model of ALM ^8^. The skewed distribution of time constants we describe is consistent with either a log-normal distribution of recurrent connections ^17^ or a diversity of connectivity profiles with graded long-range connections ^18^.

A skewed or heavy-tailed distribution of time constants has been widely observed across many brain circuits of different species ^5,2,19,20^. Notably, a similar distribution of time constants was previously shown to emerge in artificial neural networks when they are trained to solve tasks with realistically complex temporal structures ^20^. The temporal diversity in these networks appeared to promote robust and adaptive learning, suggesting that heterogeneous time constants may generally support the adaptability of learning systems in dynamic environments.

The ramping response of a single neuron is sufficient to encode time relative to a target interval. By changing the speed of this ramping, time can be encoded at variable timescales. The acceleration or deceleration of neural dynamics has previously been shown in the monkey cortex, correlating with the speed-accuracy trade-off^21^ and action timing ^22^. Our decoding results show that the range of ramping speeds in the ALM can provide timing information at varying resolutions within a fixed interval. The extent to which the speed of ramping might be adjusted in the ALM when mice encounter new temporal demands remains to be addressed in future research. We also find that the speed of ramping activity impacts the timing and stability of movement encoding in the ALM, which might affect the transmission of this information to potential downstream circuits.

What biological mechanisms might give rise to temporal heterogeneity in the ALM? One study showed that neuronal timescales across the human cortex correspond to gene expressions related to transmembrane ion transporters and NMDA and GABA receptors ^23^. Another study showed that diverse timescales across unipolar brush cells from the cerebellum are associated with the expression of metabotropic receptors, such as excitatory mGluR1 and inhibitory mGluR2/3 ^24^. Whether the expression of specific receptors can account for the diverse ramping responses in the ALM during motor planning is an interesting direction for future research.

Limitations of the present study should be considered when interpreting our results. Recently, Birnbaum et al. ^25^ showed that mice display various uninstructed movements of different body parts during the delay period of a memory-guided movement task. The dataset we analyze here does not include videos of mice performing the task, thus future work is required to determine the relationship between temporally diverse ramping dynamics and spontaneous ongoing movements. Additionally, we cannot determine whether temporal heterogeneity is required for behavior, which would require single-cell resolution perturbations ^26^.

Our findings place the ALM cortex on a long list of brain areas that display a skewed distribution of time constants. In the ALM, this diversity implies that ramping dynamics and representations of motor plans unfold over multiple timescales. From the perspective of a downstream neuron, incoming signals from the ALM cortex can provide distinct information about the direction of an upcoming movement and the remaining time until its execution. Identifying the specific input and output profiles of ALM neurons with different temporal responses is necessary to determine the implications of heterogeneous dynamics on brain functions and behavior.

## Methods

### Data description

The current paper is a secondary analysis of data previously reported in Inagaki et al. ^7^. The data was retrieved on May 15, 2020 from a publicly available database (https://crcns.org/data-sets/motor-cortex/alm-5/about-alm-5).

### Behavioral paradigm

Inagaki et al. ^7^ provided data collected from head-fixed mice engaged in an auditory-based, fixed delay task. Each trial in the task began with a series of 150 ms tones with 100 ms between each tone. The tones were either 3 or 12 kHz. This sampling period lasted 1.15 s and was immediately followed by a 2 s delay period. The delay period ended with a 100 ms auditory go cue (6 kHz carrier frequency with 360 Hz modulating frequency). The go cue indicated mice to lick the right lick-port if they were presented with 3 kHz tones during the sampling period, or the left port if they were presented with 12 kHz tones. Mice were rewarded 2 *µ*l of water if they licked the correct side within a 1.5 s response period following the go cue. Only trials when the animal correctly responded was included in the current paper.

### Electrophysiological data

The data set included processed spike trains of 755 single units that was recorded by Inagaki et al. ^7^ from the left ALM cortex of six mice across 20 sessions of auditory-based fixed delay task. Details of the recording and spike sorting procedure are described by the authors of the original report. An example MATLAB code was provided by Inagaki et al. ^7^ as part of the archived data in CRCNS to access the time stamps of spike events across the trials of each session. The authors of the original paper labeled 667 of the 755 single units as putative pyramidal neurons based on the width of their spike waveform. The procedure for identifying putative pyramidal neurons is described in detail in the original paper ^7^. Only spike trains of the putative pyramidal neurons were analyzed in the current paper. The data set also includes trials in which spikes were recorded during simultaneous photoinhibition, but we excluded these trials from the analyses in the current paper.

### Delay activity and trial-type selectivity

We used the same approach as by Inagaki et al. ^7^ to identify putative pyramidal neurons that show significant upward and downward ramping activity. A Mann-Whitney U test revealed that 516 of the 667 putative pyramidal neurons showed a significant difference between firing rate estimated within the 1 s interval prior to trial onset and firing rate estimated within the 2 s delay interval. Some of these neurons ramp up in firing during the delay (n = 302/516) while others ramp down (n = 214/516).

We estimated a normalized selectivity score for the delay activity of each neuron following the protocol used in Inagaki et al. ^7^:

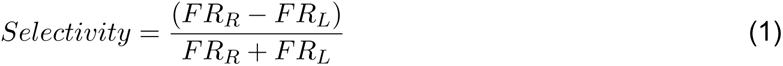

where *FR_R_* is the average firing rate (spikes/s) during the delay period across correct right-lick trials and *FR_L_* is the average firing rate (spikes/s) during the delay period across correct left-lick trials. The selectivity score ranges from -1 to 1. A positive selectivity value indicates that a neuron is more responsive during the right-lick trials and a negative value indicates that a neuron is more responsive during the left-lick trials. We used a Wilcoxon Rank Sum Test to test whether the firing rate of each neuron is different between right- and left-lick trial. The results of this test was used to determine whether each neuron is trial-type selective.

### Temporal receptive field models

To quantify the time course of activity for each neuron in the ALM cortex, we adapted an analysis method used to identify and characterize time-varying signals across monotonically firing neurons in the monkey entorhinal cortex ^5^ and stimulus-specific time cells in the monkey hippocampus and prefrontal cortex ^27^. For this analysis, temporal receptive field models were fit to the spikes of 516 putative pyramidal neurons within a 3.15 s interval preceding a Go cue, starting from the onset of the 1.15 s stimulus presentation period (t = -3.15 s relative to Go cue) to the end of the 2 s delay period (t = 0 s relative to Go cue). We considered three receptive field models: (1) a constant firing model, (2) a ramping ex-Gaussian model, and (3) a ramping ex-Gaussian model with trial-type specificity. Neurons that were identified as non-selective were fit by the second model whereas those labeled as selective were fit by the third model.

The constant model consisted of a single parameter *a*_0_ that predicted the constant probability of a spike at each time t:

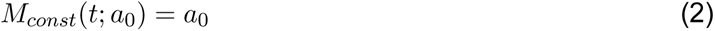

The ramping ex-Gaussian model defines the temporal modulation of the firing field as the convolution of the Gaussian function and an exponentially ramping function. The ex-Gaussian model can be numerically expressed as:

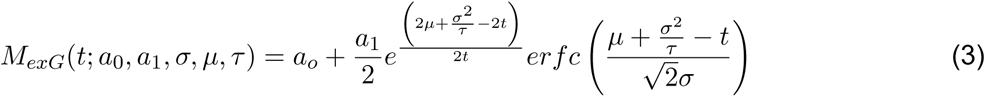

Here, *a*_0_ estimated the contribution of the constant baseline while *a*_1_ estimated the contribution of the time-varying function. A downward ramp can be achieved by letting *a*_1_ be negative. The Gaussian parameter *µ* defines the location of peak response along the time axis while *σ* determines its variability. The parameter *τ* represents the time constant of the exponential function and is defined as the time that a neuron has increased or decreased its firing 37% away from baseline. erfc is the complimentary error function.

To account for response variability across different trial types (correct, right-lick vs left-lick trials), the ramping ex-Gaussian model was extended by replacing *a*_1_ from equation 3 with two peak amplitude terms. *a*_1_ estimated the contribution of the time term for right-lick trials while *a*_2_ determined the contribution of left-lick trials. The full, trial-type specific ex-Gaussian model for the ramping time term can be numerically expressed as:

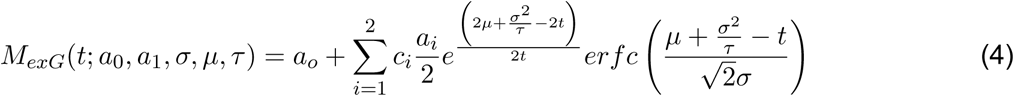

Here, *c*_1_ was set to equal 1 for right-lick trials and 0 otherwise while *c*_2_ was set to equal 1 left-lick trials only. The trial-type specific, ramping ex-Gaussian model follows the same equation, but with -t rather than t.

We calculated the fit of the receptive field models to spikes across a total of 3150 time bins (from the trial onset to the go cue) using a custom maximum likelihood estimation script run in MATLAB 2016a. The parameter *µ* was allowed to be any value between -1 s to 0 s (1 s preceding the Go cue) and *σ* was permitted to range between 0 and 0.30 s. The parameter *τ* was permitted to range between 0.01 and 5 s. The likelihood of the model fits were determined as a product of the probabilities of a spike across time bins within and across trials. We implemented the maximum likelihood parameter estimation using particle swarm (with the swarm size equal to 50). The fitting procedure was repeated until the algorithm failed to output higher likelihood for 20 consecutive iterations to avoid local minimum convergence.

We performed a goodness-of-fit test for each neuron by estimating the *R*^2^ value of the ex-Gaussian model fit to the average firing rate within the 3.15 s interval preceding the Go cue. We use this approach to exclude neurons whose model fit failed to achieve an *R*^2^ value of greater than 0.5.

### Statistical analyses

Unbiased estimates of skewness and Pearson’s kurtosis were obtained for the distribution of time constant using the *descdist* function in *R* with the argument *boot* set to 1000 for nonparametric bootstrapping. Maximum likelihood estimations were performed to fit a log-normal and gamma distribution to time constant using the *R* function *fitdist*. We reported the mean and standard error of parameter estimates for both distribution models. Log-likelihood, AIC, and BIC values were used as the goodness-of-fit measures of the fitted lognormal and gamma distributions.

### Pairwise differences

To assess the presence of diversity in time constant within and between different sessions, we measured the absolute difference between pairs of time constant values corresponding to each neuron. In the pooled condition, pairwise differences were measured between all possible pairs of neurons. In the within-session conditions, we measured time constant differences exclusively between pairs of neurons recorded within the same sessions. To measure differences in time constants across various recording sessions, we compared neurons belonging to different sessions from the same animals. These measurements were computed using a custom function in R.

### Variance component analysis

To quantify the variance in time constant values (*τ* ) attributable to differences between recording sessions and animals, we employed a generalized linear mixed-effects model (GLMM). The GLMM is formulated as follows:

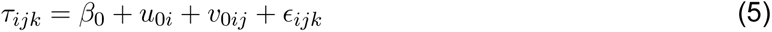

Here, *τ_ijk_* represents the time constant for the *i*-th neuron in the *k*-th session of the *j*-th animal. The term *β*_0_ is the fixed intercept, representing the overall mean time constant across all animals and sessions. The terms *u*_0*i*_ and *v*_0*ij*_ are the random intercepts for each animal and each session within an animal, respectively, and *ɛ_ijk_* is the residual error term. We assume the random effects are normally distributed with mean zero and variances 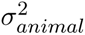 for animals and 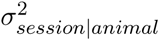 for sessions within animals. The residual variance 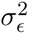 accounts for the unexplained variability in time constant. Given the right-skewed and heavy-tailed shape we observed for the distribution of time constant, we assumed *τ* to follow a gamma distribution in this GLMM. This model was implemented in R using the *glmer* function with the *family* argument set to *Gamma*. Additionally, we applied a bootstrapping approach to estimate and report the variance components relative to the total variance.

### LDA for decoding time

A linear discriminant analysis (LDA) classifier was used to decode time within the trial from the population activity of ramping neurons. The classifier was trained on even trials and tested on odd trials. We randomly sampled 125 trials for each neuron since the number of trials varied across neurons. Time within each trial was segmented into 100ms bins. Firing rate across neurons were calculated for each bin of each trial. A uniform noise, ranging from 0 to 1 *×* 10*^−^*^13^ Hz, was inserted to the firing rate in each time bin to minimize errors due to a singular covariance matrix. The averaged firing rate of each time bin for each trial across all neurons provides a sample of the training and testing data. This analysis was implemented using the MATLAB function *classify.m* using “linear” as the specified discriminant type.

### Cross-temporal decoding of trial type

To assess how movement direction, or the associated cue, is encoded by subpopulations of ramping neurons with different time constants, we trained and tested LDA classifiers to predict whether animals correctly licked left or right at every possible pair of time points throughout the trial interval ^11,7,15,12,10^. Each subpopulation of neurons comprised of 72 neurons. The range of time constant *τ* associated with each subpopulation is described in the *Results* section. Since the number of trials varied across neurons, we randomly sampled 125 trials for each neuron in slow-, medium-, and fast-ramping subpopulation I, II, and III. We segmented the 2 s delay intervals into 100 ms bins. We trained the classifiers on 80% of the sampled trials and tested them on the remaining 20% of the trials for each pair of time bins. The firing activity across neurons were calculated for each bin of each trial. We used the MATLAB function *rref.m* to reduce the training data to full rank and identify the pivots before each run of the classifier. This was performed to ensure the stability of the LDA. The LDA classifier was then implemented using the MATLAB function *classify.m*. We repeated training and testing procedures for 20 iterations. To obtain robust results, we randomly sampled subsets of trials and 71 of 72 neurons in each subpopulation at each iteration. This procedure yielded an accuracy matrix for each of the three subpopulations. The matrix is composed of pairs of training and testing time bins. Identical training and testing time bins lie along the diagonal of this matrix. The off-diagonal values of the accuracy matrix indicate the similarity in how information is encoded by the population between two different time points.

### Alternative and independent estimation of time constants

We used the Zeraati et al. ^6^ model to independently evaluate time constants for neurons designated as either ramp or decay. Their unbiased method uses a generative model to evaluate the time constants from the sample autocorrelation of an Ornstein–Uhlenbeck (OU) process as opposed to the biased method of fitting an exponential curve directly to the autocorrelation function of a time series. Using their provided abcTau python package, we fit a one tau OU process model and a two tau OU process model to the average firing rates of each neuron across a 3150 ms interval preceding the Go cue. We then estimated the autocorrelation of each neuron with a binsize of 50 ms. For the one tau OU process, we assumed a uniform prior with the min tau = 0 ms and max tau = 10000 ms. For the two tau OU process, we assumed the same distribution for *τ*_1_ and a uniform prior with the min tau = 100 ms and max tau = 10000 ms for *τ*_2_, with *τ*_2_ *> τ*_1_ enforced. The prior for coefficient or weight for the first timescale was uniform between 0 and 1. We used a linear distance function and model hyperparameters *epsilon*_0_ = 1, *minsamples* = 100, *steps* = 60, and *minAccRate* = 0.01. We then used the inbuilt model comparison *abcTau.compcdf()* module to calculate the Bayes factor between the two models for each cell. If the one tau OU model fit better, that tau was taken as the timescale for that cell. Otherwise, *τ*_2_ from the two tau OU process was taken as the timescale for that cell.

## Acknowledgments

We thank Hidehiko Inagaki and colleagues from the Svoboda Lab for sharing their data on the Collaborative Research in Computational Neuroscience (CRCNS) website. We thank Mike Economo for helpful comments and ideas on the paper. We thank Spencer Byers and Tereza Dafalia for valuable feedback on the manuscript. ROA was supported by a NSF GRFP Grant No. DGE-1840990 and NIH Blueprint DSPAN Award no. F99NS130925.

## Author Contributions

Conceptualization: R.O.A. and M.W.H.; modeling and analysis: R.O.A., I.M.B., N.A.C., and L.N.P.; writing – original draft: R.O.A., L.N.P., and M.W.H.; writing – review & editing: R.O.A., M.W.H., and B.B.S. This work was completed equally in the labs of M.W.H. and B.B.S.

## Supplementary Materials

**Figure S1:**
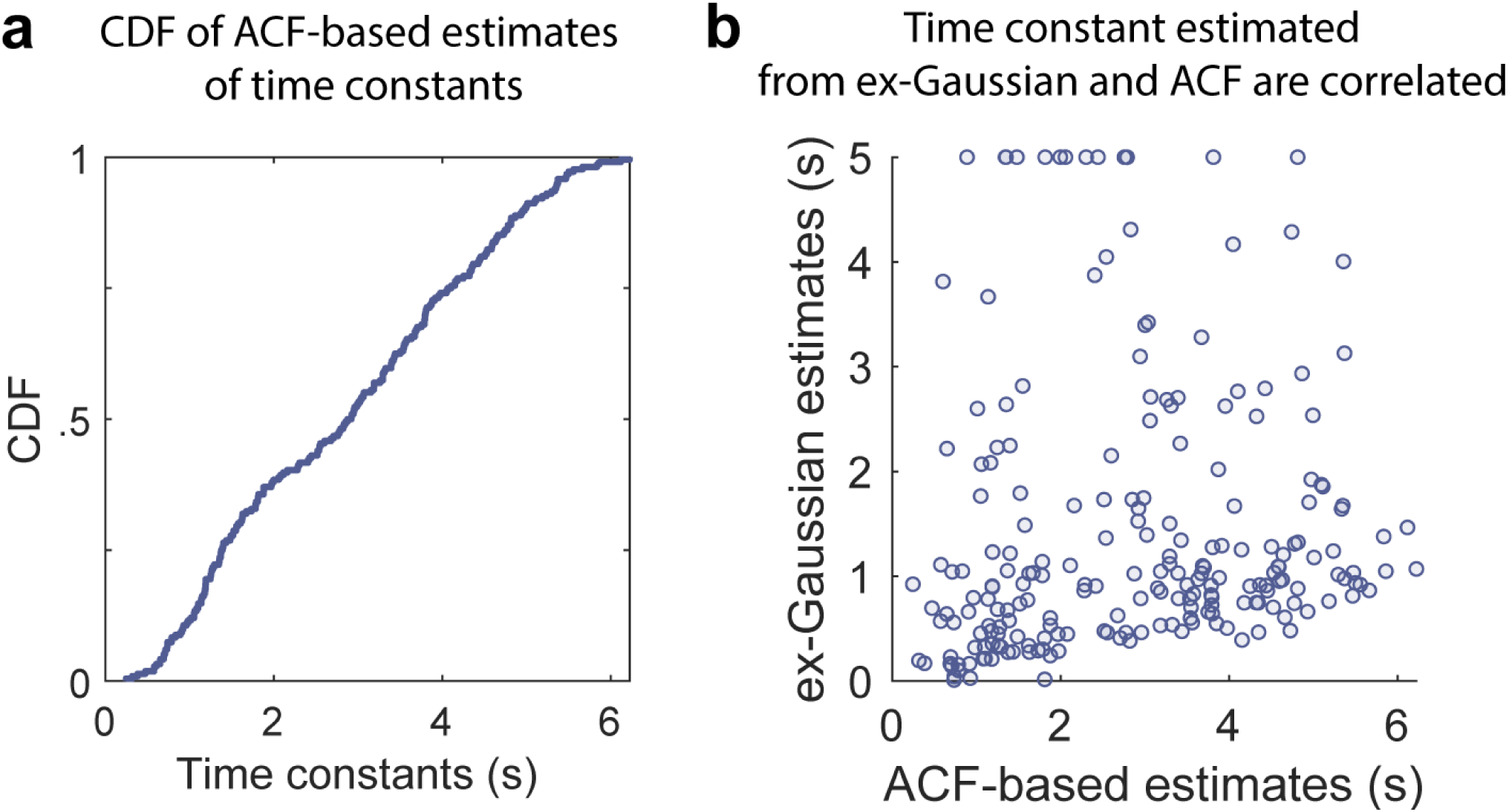
Alternative estimation of time constants of ramp cells. **a.** Cumulative distribution of time constants (median = 2.90 s, interquartile range = 1.39 s to 4.07 s) estimated from the autocorrelation function (ACF) of neurons using the approach previously reported by citetzeraati2022flexible. **b.** Scatterplot of time constants estimated from the ex-Gaussian model and the ACF-based approach. A significant correlation between the two sets of estimates suggests consistency in time constant measurements across the two methods (Kendall’s *τ* = 0.19, *p < .*001).

**Figure S2:**
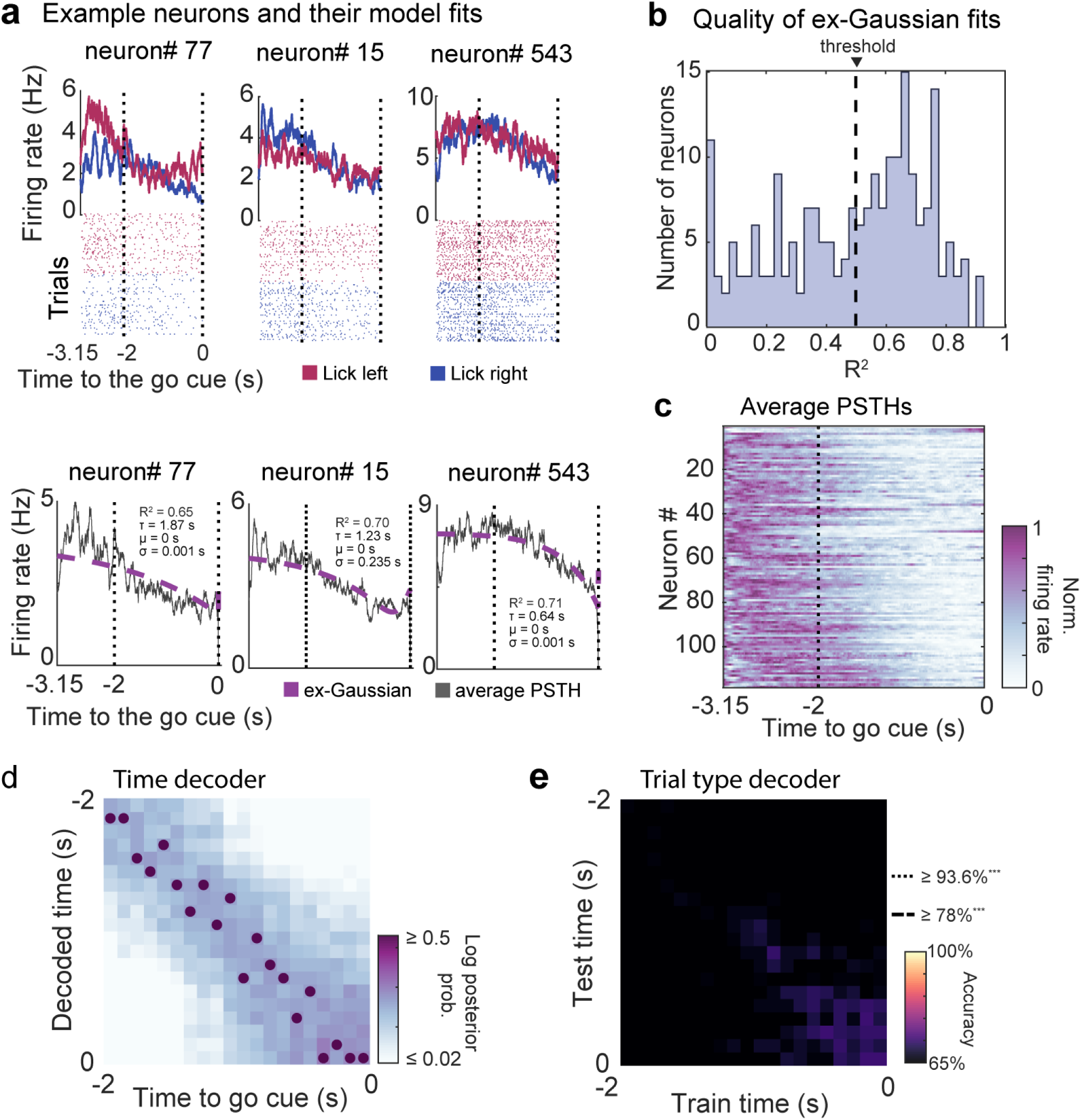
Downward ramping neurons. **a.** Firing rate and rasters of three example neurons with downward ramping activity. Blue, low-pitch tone and lick-right. Red, high-pitch tone and lick-left. In the bottom row are the ex-Gaussian model fits. **b.** Quality of model fit for downward ramping neurons. A proportion of downward ramping neurons were fit by the ex-Gaussian model above threshold (dashed line). **c.** Heatmap of the average PSTHs of downward ramping neurons. **d.** LDA matrix from the decoding of time using the population activity of downward ramping neurons. **e.** Cross-temporal decoding of trial type from the population activity of downward ramping neurons

